# Microbiota of the alien species *Paraleucilla magna* (Porifera, Calcarea) from the Southwestern Atlantic, and a comparison with that of other calcareous sponges

**DOI:** 10.1101/626192

**Authors:** Maira Ferreira Lopes, Beatriz Mágna, Michelle Klautau, Eduardo Leal Esteves, Rodolpho M. Albano

**Author notes:** Corresponding author: Rodolpho M. Albano.

## Abstract

Sponges (Porifera) co-evolved with microorganisms in a well-established symbiotic relationship. Based on this characteristic, sponges can be separated into high microbial abundance (HMA) and low microbial abundance (LMA) species. *Paraleucilla magna* (Calcarea, Porifera) is an alien species of ecological importance in the Brazilian coastline. Little is known about the composition of its microbiota and that of other calcareous species, especially those inhabiting the Southwest Atlantic. Here, we describe the microbiota of *P. magna* and compare it to that of other calcareous sponge species for which such data exist. *P. magna*’s microbiota shows a lower diversity than that of *Clathrina clathrus*, *C. coriacea*, *Leucosolenia* sp., *Leuconia* sp. and *Leucetta antarctica*. *P. magna* microbiota is dominated by two bacterial OTUs of the Alphaproteobacteria class, that could not be classified beyond class (OTU001) and family levels (OTU002; Rhodospirillaceae). The Thaumarcheota was the predominant archaeal phylum in *P. magna*, with OTUs mainly affiliated to the genus candidatus *Nitrosopumilus*. The comparison with other calcareous species showed that microbiota composition correlated well with sponge phylogenetic affiliation. Metabolic prediction with PICRUSt software of *P. magna* bacterial microbiota indicated that membrane transport and carbohydrate, amino acid and energy metabolisms were most abundant while, for the archaeal domain, pathways related to translation, and energy metabolisms were predominant. Predicted metabolic features were compared between the different sponge species and seawater samples, showing that pathways related to cell motility, membrane transport, genetic information processing, xenobiotics metabolism and signal transduction are higher in the former while amino acid and nucleotide metabolism, translation, replication and repair, folding, sorting and degradation and glycan biosynthesis and metabolism are abundant in the latter. This study shows that *P. magna*’s microbiota is typical of an LMA sponge and that it differs from the microbiota of other calcareous sponges both in its composition and in predicted metabolic pathways.

## Introduction

Sponges (phylum Porifera) represent the most ancient metazoan lineage. For over 600 million years, their presence has been recorded in a great variety of habitats while, at the same time, maintaining a simple bauplan. Several studies point to the conclusion that the key to this adaptability may lie on their long association with microbes [1].

Sponges co-evolved with microorganisms forming, through time, a well-established symbiotic relationship. While the host provides nutrition from the degradation of other microbes and a habitat in its mesohyl, the prokaryotic community enhances its metabolic potential with complementary physiological functions and the production of useful secondary metabolites [2].

In the last two decades, advancements in next generation sequencing (NGS), in bioinformatics tools and the growth of gene marker databases, allowed a deeper study of this relationship. Nowadays, for example, it is known that organisms from the three domains of life inhabit sponges, with different levels of specificity [2,3]. The nature of this symbiotic relationship can be used to divide sponges into two main categories: high microbial abundance (HMA), in which a diverse microbiota can account for up to 35% of the host’s biomass and low microbial abundance (LMA), comprising sponges that are inhabited by fewer and less diverse microbes. In the latter, the most common scenario is the predominance of only one or two microbial phyla [4].

Another great motivation for gaining a better understanding of this relationship is the potential biotechnological applications of enzymes and bioactive molecules produced by microbial sponge symbionts. As reported by Abdelmohsen *et al.* [5], 57% of the marine natural products with described action against drug resistant pathogens (parasites, fungi and viruses) were extracted from species of the phylum Porifera.

Porifera has four extant classes that are characterized by different kinds of skeletal features: Demospongiae, Calcarea, Hexactinellida and Homoscleromorpha. The composition and dynamics of microbial symbionts and how they relate to taxonomic ranks in Demospongiae have been extensively studied [6]. This concentration of resources in the class comprising the majority of Porifera’s described species has left a deficit of information on the microbiota of species belonging to the other classes. Additionally, there is still limited data on the microbiota of sponges of the South Atlantic. To date, few studies have analyzed the microbiota of calcareous sponges using high-throughput technology and the only species for which such data exist are *Clathrina clathrus*, *C. coriacea*, *Leucosolenia* sp., *Leuconia* sp. and *Leucetta antarctica* [7,8].

In the present study, we aim to shorten this gap by describing the microbial symbionts (Bacteria and Archaea) of *Paraleucilla magna* Klautau, Monteiro & Borojević, 2004 (Porifera, Calcarea) using next generation sequencing. This species was first described in the 1980’s in Rio de Janeiro state [9] where it has been considered a cryptogenic species. Further records in Brazil were in São Paulo and Santa Catarina states, but *P. magna* has also been registered in several localities in the Mediterranean Sea [10–12], and in the Eastern Atlantic Ocean [13]. *P. magna* harbors a variety of organisms, such as crustaceans, molluscs, bryozoans, and polychaetes [14] and some bacteria that produce antimicrobial compounds have also been isolated from tissue samples of this species [15].

*P. magna* is capable of reproducing throughout the year and produces high numbers of larvae [16], two characteristics that indicate an efficient invasive potential [17]. Populations of *P. magna* present high demographic fluctuations and have not presented any visible harm to native Brazilian sponge populations. It is also one of the most common calcareous sponge species on the rocky coasts of Rio de Janeiro. Moreover, in the Mediterranean Sea it is considered an invasive species, as it causes problems in mollusc farms.

To increase our knowledge about the symbionts of Brazilian sponges we analyzed the microbiota and the predicted metagenome of *P. magna* collected at Marica’s archipelago in Rio de Janeiro, Brazil and performed a comparative analysis with the microbiota of other calcareous species with available data.

## Material and Methods

### Sample collection and storage

*Paraleucilla magna* individuals (n=2) that were at least 10 meters apart were collected by scuba diving at a depth of 6 meters in Maricás archipelago, Rio de Janeiro (23°00’41.8’’ S, 42°55’07.39’’ W), Brazil, in February 2013 (Fig 1). Excess seawater was removed by gently pressing the samples against sterile filter paper. A fragment of approximately 500 mg was excised from each individual with a sterile scalpel blade. Fragments were stored in RNAlater (Qiagen) at room temperature and transported to the laboratory for DNA extraction. A voucher of each specimen was deposited at the Porifera Collection of the Rio de Janeiro State University (UERJPOR) under voucher numbers UERJPOR27 (*P. magna* 6) and UERJPOR26 (*P. magna* 8), respectively. Morphological identification was performed by three specialists, BM, MK and ELE. For microbiota analysis of the surrounding seawater, 5 liters of surface seawater were collected in a sterile container and filtered through a 0.22 µm membrane (Millipore). The membrane was stored at −20 °C until processing.

**Figure 1.**
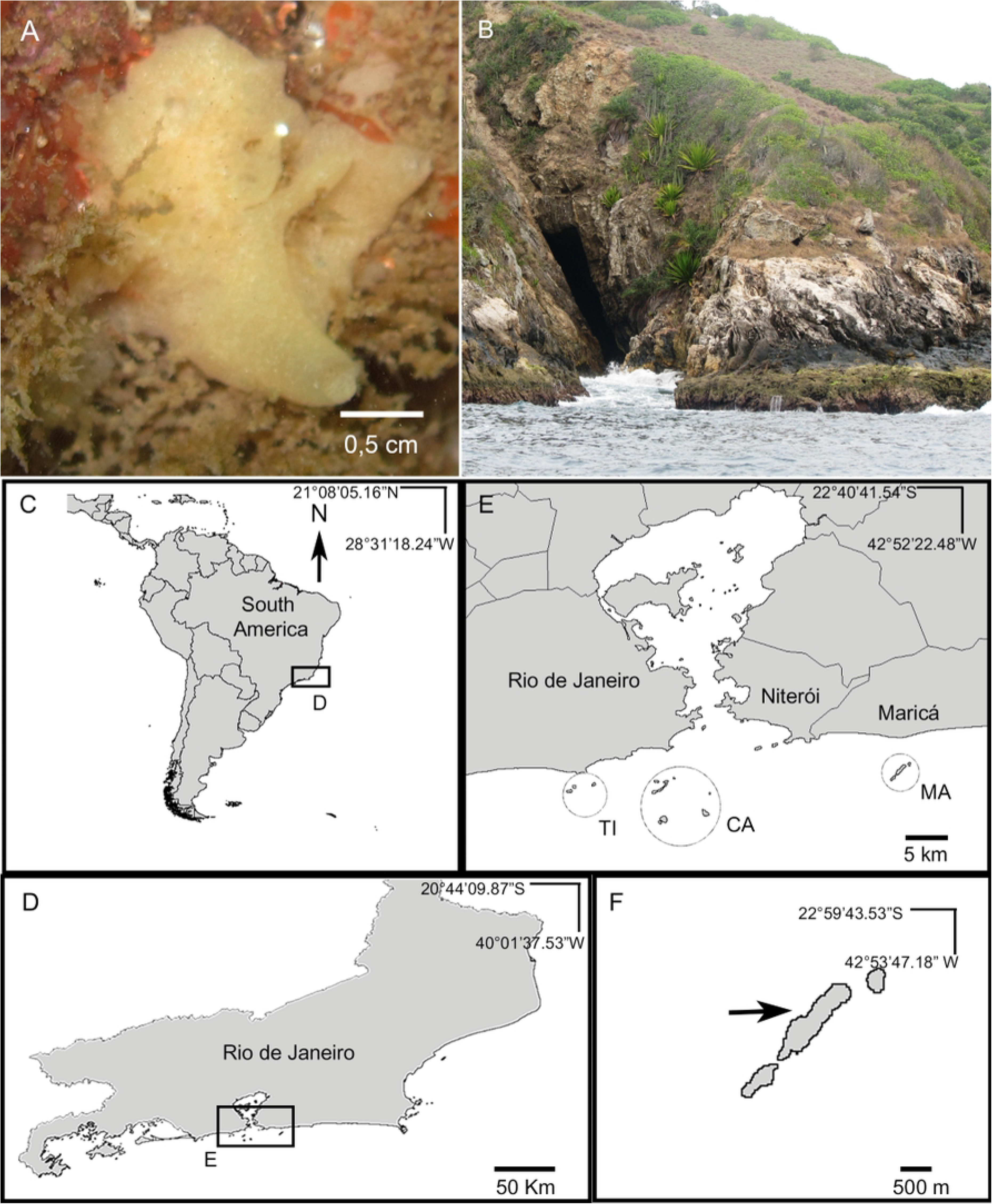
*P. magna in situ* photograph and sample collection locality. (A) *In situ* photograph of one of the *P. magna* individuals collected for microbiota analyses (*P. magna* 6). (B) Main island of Maricás Archipelago showing a vertical rocky slope, a typical environment in this island. (C) Map of South America showing Rio de Janeiro state in the inset. (D) Map of Rio de Janeiro state with collection area in the inset. (E) Central coast of Rio de Janeiro state including Guanabara Bay and nearby islands: Cagarras Archipelago (CA), Maricás Archipelago (MA) and Tijucas Archipelago (TI). (F) Collection locality in Maricás Archipelago.

### DNA extraction

For more efficient extraction, small pieces of sponge tissue were cut with a sterile scalpel and macerated with sterile polypropylene pestles in microcentrifuge tubes containing extraction buffer. The filter membrane was cut into small pieces with sterilized scissors and placed into a microcentrifuge tube with the same buffer. Total DNA extractions were carried out with the DNEasy blood and tissue kit (Qiagen) following the manufacturer’s recommendations. DNA quantification was performed using the Qubit High sensitivity ds DNA assay (Thermo Fisher Scientific, Brazil).

### Sponge phylogenetic analysis

To provide a molecular tag for our *P. magna* samples, amplification of the 18S rRNA gene was performed as described by Redmond *et al.* (2007)[18]. The amplification products were purified, sequenced by Sanger methodology with the BigDye reagent (Thermofisher, São Paulo, Brazil) and analyzed on an Applied Biosystems 3500 Genetic Analyzer capillary instrument. A minimum of three sequencing reactions was performed for each fragment on both forward and reverse strands. Contigs from FASTA formatted sequences were constructed with Geneious R9 software (Biomatters) and used for BLAST [19] searches against GenBank [20]. The resulting sequences were deposited under accession numbers KY634245 and KY634237.

### Marker gene library construction and sequencing

For the analysis of the microbiota, the V4 region of the 16S rRNA gene was amplified using primers and PCR conditions as described by Caporaso *et al*. (2011)[23]. Triplicate PCR reactions for each sample were pooled, purified, quantified with the Qubit fluorometer and paired-end sequenced with the 500 cycles MiSeq reagent kit V2 on a MiSeq instrument (Illumina). Sequences were deposited on SRA databank under accession number SRP127694.

### Processing of the sequence reads and microbiota analysis

The sequence reads were processed with mothur software (v 1.39.5)[24] according to the Miseq SOP [25]. Briefly, paired-end reads were joined using the make.contigs command and filtered to exclude ambiguities (max=0), homopolymers (maxhomop=8), sequences over 252 bp and to ensure correct overlapping. Then, they were aligned against the Silva (v. 128) database [26], pre-clustered (diffs=2) and screened for chimeras, using the VSEARCH algorithm [27]. Screened sequences were classified, and those belonging to undesired taxa such as chloroplast, mitochondria and Eukarya were removed. Analysis was carried out separately for Bacteria and Archaea domains. To build operational taxonomic units (OTUs), clustering was set at 97% similarity.

For taxonomic description, sponge OTUs with relative abundance lower than 1% of the total number of sequences were grouped as others; this cutoff was defined at 5% for seawater OTUs. Rarefaction curves and alpha and beta diversity analysis were calculated after subsampling groups to the smallest library size. Results were plotted with Prism v7 software (GraphPad, USA).

A more thorough investigation was performed on two OTUs (OTU001 and OTU002) that presented a high relative abundance in *P. magna*. The get.oturep command was used within mothur software to obtain the representative fasta sequences for both OTUs. These sequences were aligned against sequences from a *P. magna* 16S rRNA gene clone library that was constructed by amplifying this gene from genomic DNA with primers 27F (5′-AGAGTTTGATCCTGGCTCAG-3′) and 1492R (5′-GGTTACCTTGTTACGACTT-3′). The PCR amplification products were cloned into the PGEMT vector (Promega) and inserted into *E. coli* XL1Blue cells. Transformed bacterial colonies were picked, plasmid DNA was recovered and used in sequencing reactions to obtain full-length 16S rRNA gene sequences.

Sequences that presented >97% identity were chosen as full-length counterparts of the OTU sequences and were used on BLAST searches against the GenBank database. The best-hit sequences were downloaded and aligned with the OTUs’ representative sequences using MAFFT algorithm. A maximum likelihood tree was built using the GTR model/ G+I and 500 bootstraps as a test of phylogeny with MEGA v7 software [22].

### Comparison with the microbiota of other calcareous sponges

Raw V4 sequence reads of 16S rRNA surveys from the calcareous sponge species *Clathrina clathrus*, *C. coriacea*, *Leucosolenia* sp., *Leuconia* sp. and *Leucetta antarctica* and from seawater samples from the Antarctic region, Spain and Curaçao that are available at SRA were downloaded (Supplementary material). These sequences were analyzed along with our data set within mothur software essentially as described above. However, the maximum length parameter was altered to 125 bp to match the other samples’ length. As there were multiple biological replicates from each sample, those replicates that did not reach 90% coverage were removed.

A biom formatted OTU table was used to calculate and plot alpha diversity within the Microbiome Analyst website (http://www.microbiomeanalyst.ca/) [28]. The following parameters were used: The low count filter was set without a minimum count and the prevalence in samples was set at 10%. The low variance filter was set at inter-quartile range and the percentage to remove was set at 0%. The data was then normalized by rarefying to the minimum library size and scaled by total sum scaling.

To determine the similarities of the microbial communities among the different sponge and seawater samples a dendrogram was constructed with mothur from Bray-Curtis distances obtained from samples rarefied to the minimum library size. Weighted unifrac was the chosen distance metric for beta diversity analysis from the rarefied samples, which was visualized on a NMDS graph.

### Functional category prediction with PICRUSt software

For metagenome prediction, sequences were processed within mothur as outlined above but, instead of the Silva database, the Greengenes database (gg_13_5_99) [29] was used to align and classify the OTUs. Three biom formatted files were produced with the make.biom command in mothur containing OTU counts and Greengenes taxonomy labels in the PICRUSt format: Two with OTUs from the *P. magna* samples and the surrounding seawater of Maricás archipelago (one for the Bacteria and one for the Archaea domain), and a third one for the bacterial domains of all calcareous sponge and seawater samples.

These biom formated files were uploaded to a Galaxy web application maintained by the Huttenhower lab (https://huttenhower.sph.harvard.edu/galaxy/). Using PICRUSt software[30], OTU counts were normalized by 16S copy number and the metagenome was predicted in the form of KEGG ortholog abundances [31] that was used for functional categorization.

The PICRUSt output in biom file format was uploaded for analysis in the Calypso online software (http://cgenome.net/wiki/index.php/Calypso; [32]) where the 20 most abundant features were presented in heatmaps for *P. magna* samples description. For the comparison with other sponge species, the 15 most abundant features were chosen. Similarities in the predicted metagenomes among all calcareous sponges and seawater samples were determined with a hierarchical clustering graph according to Bray-Curtis distances that was also computed within the Calypso platform. The parameters used for these analyses on the Calypso platform were the following: Data was filtered by removing rare taxa with less than 1% relative abundance, and the data were normalized by total sum normalization followed by square root transformation. Statistical analysis was performed in STAMP [33], using two groups comparison between Sponges and Seawater, using the two-sided Welch’s t-test with Benjamini-Hochberg FDR correction.

## Results

### Sequencing run details and alpha diversity

Sequencing of the V4 tag of the 16S rRNA gene produced, after mothur processing, 819,538 sequence reads for the two *P. magna* individuals, being 98.5% bacterial and 1.5% archaeal (Table 1). From the seawater sample of Maricás archipelago, 216,345 sequences were obtained, 93% bacterial and 7% archaeal. These sequences resulted on a total of 2,214 and 2,034 bacterial OTUs obtained for *P. magna* 6 and 8, respectively, while 2,576 OTUs were retrieved for the seawater sample. We recovered fewer archaeal OTUs which were 24, 16 and 55 for *P. magna* 6, *P. magna* 8 and seawater samples, respectively.

**Table 1.** Sequence data and alpha diversity. Sobs: species observed; Invsimpson: Inverse Simpson index.

| Sample | Taxon | Nseqs | Coverage | Sobs | Invsimpson |
| --- | --- | --- | --- | --- | --- |
| *P. magna* 6 | Bacteria | 323,510 | 0.998 | 2214 | 2.98 |
| *P. magna* 8 | Bacteria | 483,744 | 0.999 | 2034 | 2.47 |
| Seawater | Bacteria | 201,156 | 0.995 | 2576 | 24.00 |
| *P. magna* 6 | Archaea | 8,569 | 0.999 | 24 | 1.23 |
| *P. magna* 8 | Archaea | 3,715 | 0.999 | 16 | 1.26 |
| Seawater | Archaea | 15,189 | 0.999 | 55 | 4.72 |

Although the species observed index (Sobs) presented little difference between seawater and sponge samples, a higher diversity was found in seawater, as indicated by the inverse Simpson (invSimpson) index, which measures richness as well as evenness (Table 1, Fig S1). All libraries had high coverage, above 99%, and reached a plateau in rarefaction curves (Fig S1).

### Taxonomic composition of *P. magna* and surrounding seawater microbiota

The bacterial communities of both *P. magna* individuals and the surrounding seawater are composed of few phyla with relative abundance above 1% or 5%, for sponges and seawater samples, respectively. Proteobacteria is the most abundant, with the class Alphaproteobacteria representing 84%, 92% and 21% of the OTUs in *P. magna* 6, 8 and seawater, respectively (Fig 2). Gammaproteobacteria was more prominent in seawater sample, represented by 43% of the OTUs, while in *P. magna* 6 and 8 only 5% and 4% of them, respectively, were affiliated with this taxon. Bacteroidetes was the second most abundant phylum, especially in seawater in which it amounted to 30% of the OTUs, while in *P. magna* 6 and 8 the phylum’s proportion was 3% and 2%, respectively. Planctomycetes was present only in the seawater sample (2%) and *P. magna* 6 (1%). OTUs affiliated with the phylum Marinimicrobia comprised 1% of the total in seawater (Fig 2).

**Figure 2.**
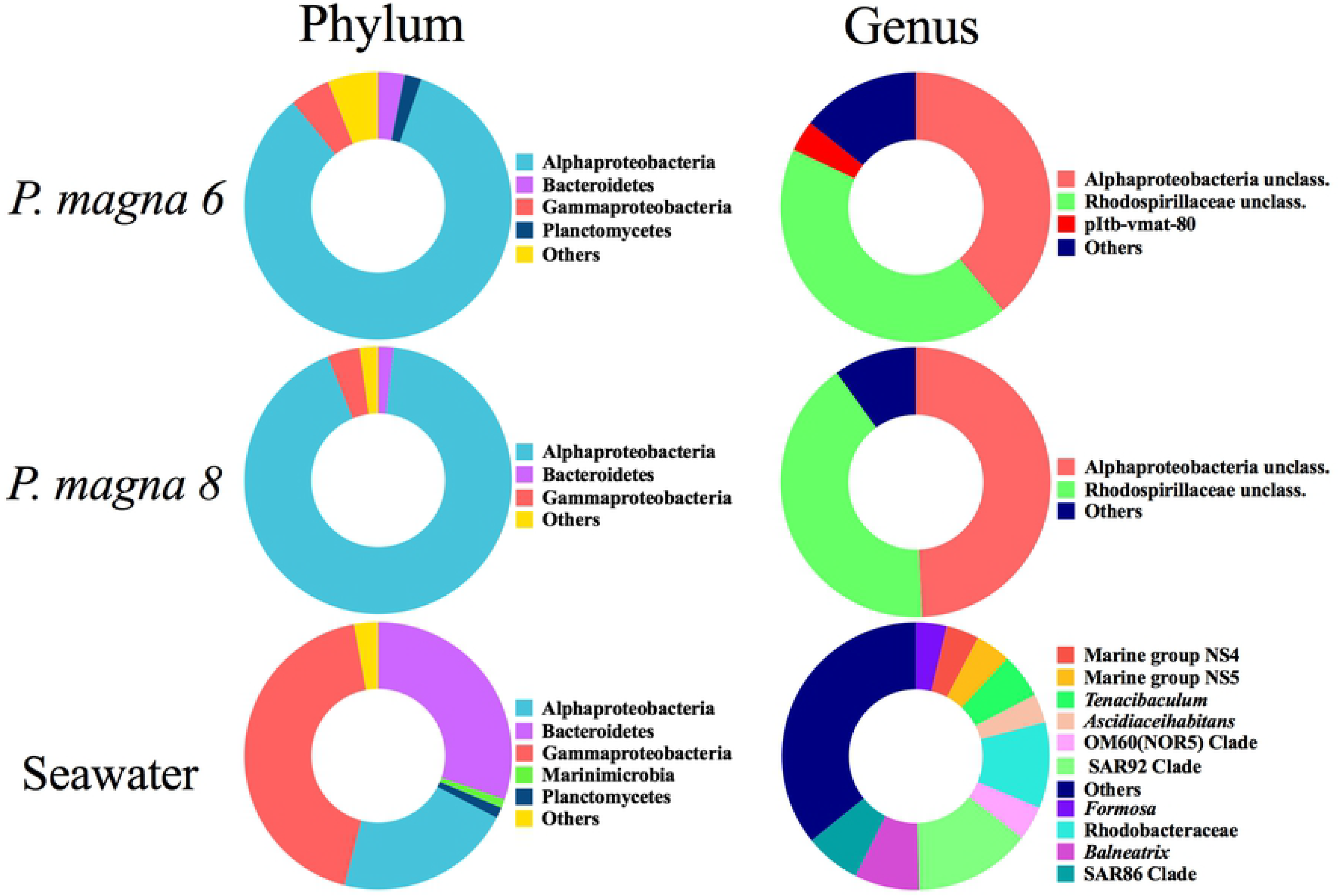
Taxonomic composition of the Bacteria domain from the microbiota of *P. magna* and seawater samples. Only taxa with relative abundance above 1% (for sponges) or 5% (seawater) are shown. Others represents taxa below these cutoffs. The predominant phylum Proteobacteria was separated into classes.

At a lower taxonomic rank, both *P. magna* individuals presented two Alphaproteobacteria OTUs, hereafter named OTU001 and OTU002, as dominant taxa. OTU001 has a relative abundance of 38% and 49% in *P. magna* 6 and 8, respectively, and could not be classified beyond the class level. When its representative sequence was used for searches against full-length sequences obtained from a 16S rRNA clone library, a clone (A3) with 98% identity was found. The Blast search of this sequence against GenBank database retrieved sequences with at most 91% similarity. These sequences originated from uncultured bacteria, mostly belonging to the class Alphaproteobacteria; those were isolated from diverse marine environments. When these sequences were used in a phylogenetic analysis, OTU001 and its full-length representative sequence formed a separate cluster with strong bootstrap support indicating that they might belong to a novel species or, perhaps, genus with low representation in databases (Fig 3A).

**Figure 3.**
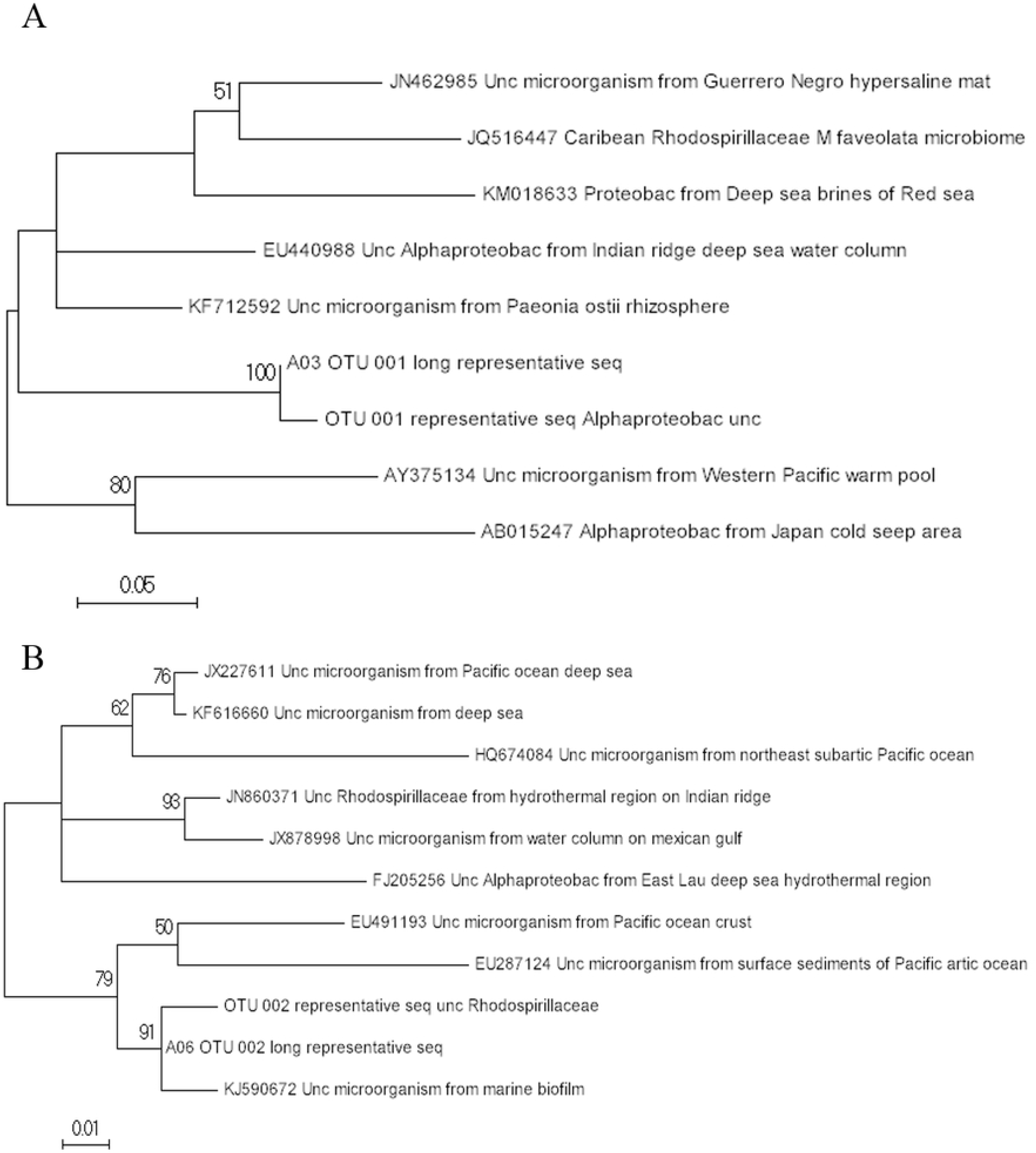
Phylogenetic analysis of *P. magna*’s predominant bacterial OTUs. Maximum lilkelihood trees of full length 16S rRNA gene sequences were constructed with Mega software. (A) OTU001 and closest relatives retrieved from Blast searches against Genbank. (B) OTU002 and closest relatives retrieved from Blast searches against Genbank. Only bootstrap values above 50% are shown.

OTU002 is a member of the Rhodospirillaceae family, which could not be classified to the genus level. It has a relative abundance of 43% and 41% in *P. magna*’s 6 and 8 microbiota, respectively. As described above, a representative sequence was aligned against sequences from our clone library, and clone A6 showed 99% identity with the V4 region of OTU002. The full-length sequence was used on a Blast search against GenBank and sequences from Alphaproteobacteria found in marine environments were retrieved showing identities up to 95%. Upon a phylogenetic analysis (Fig 3B), our sequences grouped closely with bacteria isolated mainly from coastal and intertidal regions. So, although no further taxonomic information could be obtained, we could infer that this sequence belongs to an already reported, unnamed genus of marine bacteria.

An OTU affiliated with pItb-vmat-80, a genus of environmental Gammaproteobacteria isolated from a microbial mat found in a shallow submarine hot spring off the coast of Japan [34], was also observed only in *P. magna* 6.

OTUs found in the seawater sample were affiliated with a great diversity of genera, and 35% of them did not reach a minimum relative abundance of 5%. Typical marine genera were also observed such as those affiliated with the SAR and NOR clades, with SAR_92 clade being the most abundant, with 14% of the OTUs (Fig 2).

### Taxonomic composition of the archaeal domain

Two archaeal phyla were found in all samples with variable relative abundance. While sponge OTUs were mostly affiliated with the phylum Thaumarcheota (99% and 96% for *P. magna* 6 and 8, respectively), seawater sample OTUs were placed in nearly equal proportion within the phyla Thaumarcheota (47%) and Euryarcheota (53%). At the genus level, a greater difference between the sponge and seawater samples was observed. In *P. magna* individuals, the candidatus *Nitrosopumilus* was dominant (91% and 89% in *P. magna* 6 and 8, respectively) with a smaller contribution of candidatus *Nitrosopelagicus* and of subtypes Marine Group I and II. Meanwhile, in the seawater library half of the OTUs were affiliated with Marine Group II (51%), followed by those related to candidatus *Nitrosopumilus* (35%) and *Nitrosopelagicus* (11%) (Fig 4).

**Figure 4.**
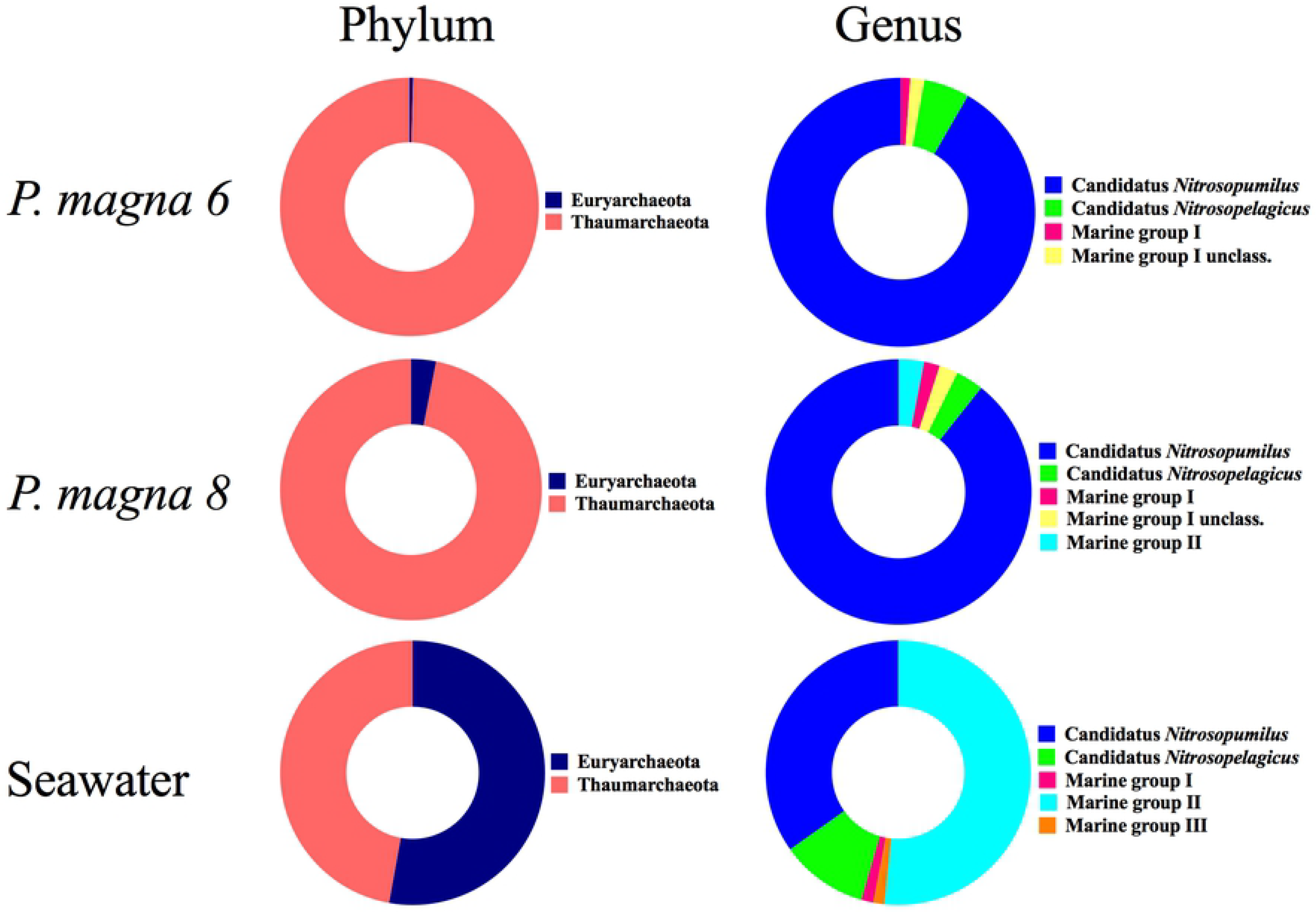
Taxonomic composition of the Archaea domain from the microbiota of P. magna and seawater samples. Only taxa with relative abundance above 1% are shown.

### Functional category prediction of the bacterial and archaeal microbiomes

PICRUSt software was used to predict metabolic pathways present in the microbiomes of *P. magna* and surrounding seawater. A heatmap of the 20 most abundant KEGG pathways at hierarchical level 2 shows that the functional profiles were clearly clustered according to their source for both bacterial and archaeal microbiomes (Figs 5A and B). The most abundant pathways in the *P. magna* bacterial communities are those related to membrane transport and carbohydrate, amino acid and energy metabolism, indicating the predominance of a heterotrophic metabolism. Replication and repair, nucleotide metabolism and translation pathways were higher in seawater bacterial communities, indicative of a higher proliferation potential in the open sea. Signal transduction, metabolism of other amino acids, terpenoids and polyketides, folding, sorting and degradation, general metabolism and transcription pathways were the least represented in all samples (Fig 5A).

**Figure 5.**
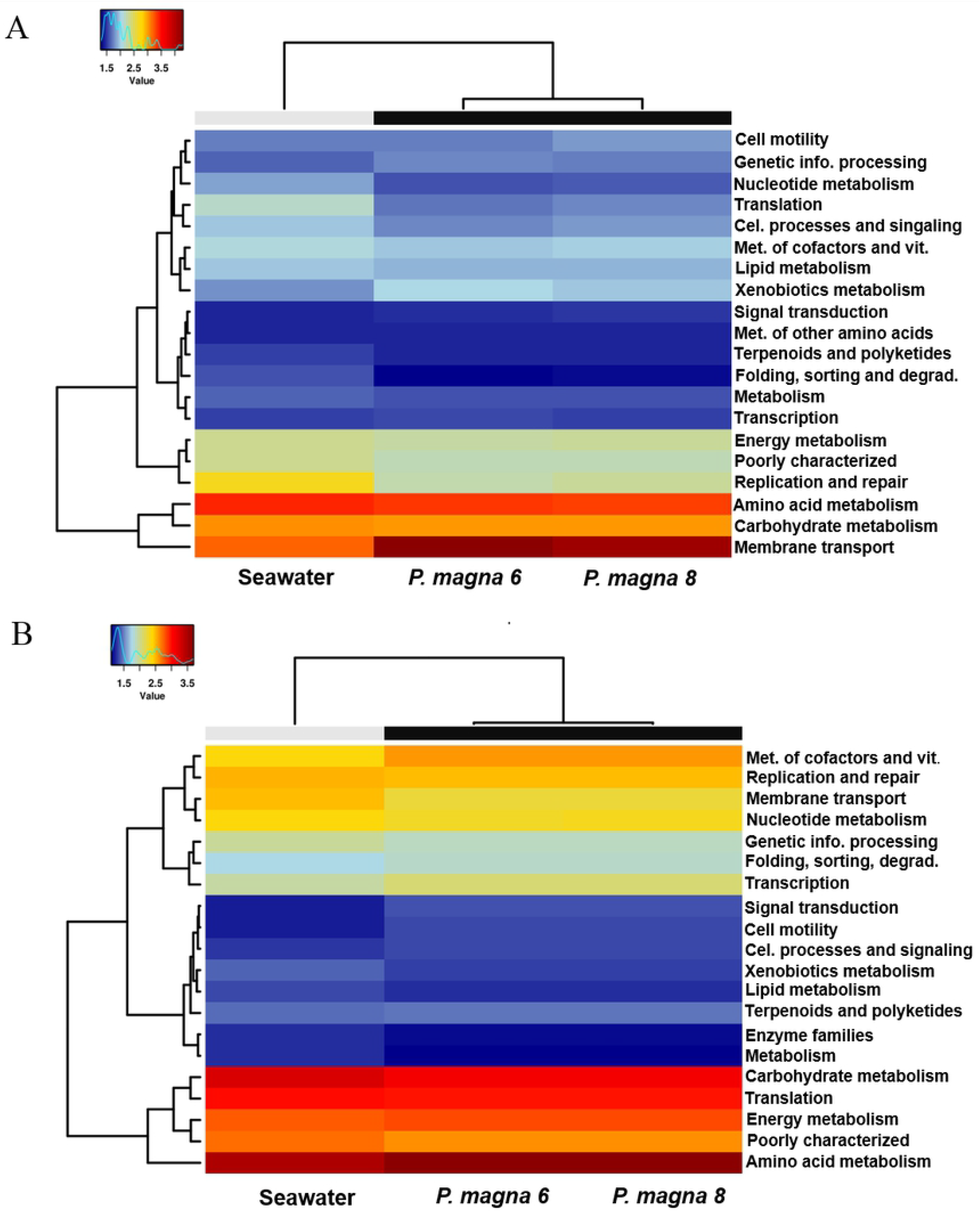
Abundance heatmap of predicted metagenomic functional profiles from *P. magna* and seawater samples. The 20 most abundant metabolic pathways predicted with PICRUSt for Bacteria (A), and Archaea domains (B) are shown. Colors shift from blue (lower) to red (higher) according to pathway abundance in each sample.

The predicted metagenome of archaeal communities showed that pathways related to translation, and energy metabolism, the latter involving amino acids and carbohydrate metabolism, were the most abundant. Among these pathways, only carbohydrate metabolism was more prominent in seawater. Differences were found in intermediately abundant pathways, where metabolism of cofactors and vitamins were more prevalent in sponges while replication and repair and membrane transport were more prominent in seawater. Among features with smallest abundance, enzyme families and metabolism were lower in the sponges and signal transduction, cell motility, processes and signaling pathways were lower in seawater (Fig 5B).

### Comparative analysis of the microbiota of different calcareous sponge species

In order to compare the composition of the microbiota present in our samples and in other calcareous sponge species, we processed all datasets together within mothur software. Weighted unifrac phylogenetic distances were plotted on an NMDS graph in which we observed that, as expected, *P. magna* samples clustered together (Fig 6). The clustering of biological replicates of the same species is also observed, indicating stability in the core microbiota. *Clathrina clathrus* and *C. coriacea*, species of the same genus, had the most similar communities, indicating that phylogenetic distance and microbiota membership are associated. The microbiota from the seawater samples show more similarity in composition among themselves than with the sponge samples and, therefore, are closer to each other in the plot. Similar results can be seen on a cladogram obtained using a Yue & Clayton distance matrix where seawater and sponge samples were also separated, and community similarity mirrored taxonomic affiliation (Fig S2).

**Figure 6.**
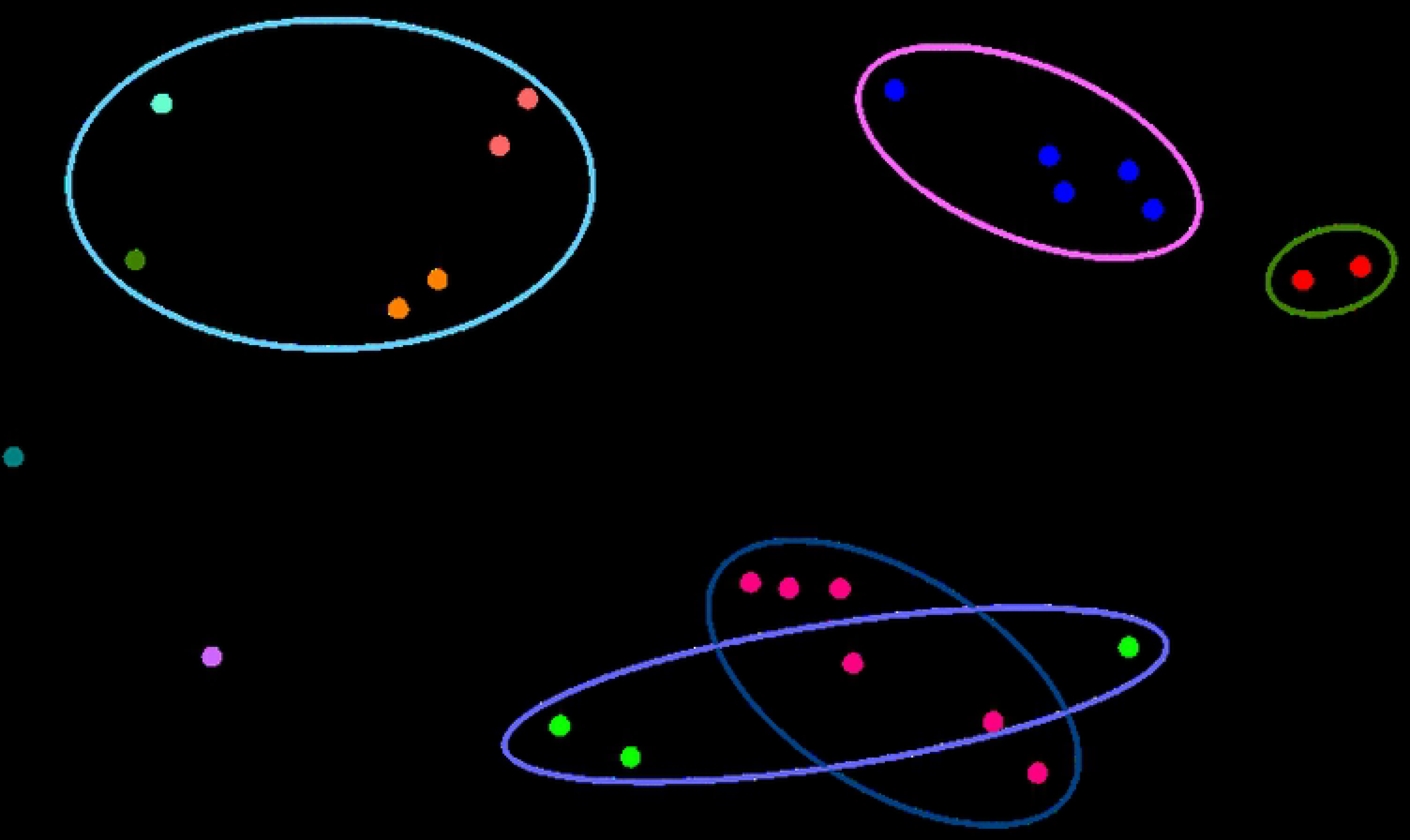
Beta diversity analysis of microbiota composition from different calcareous sponge species and seawater samples. Weighted unifrac distances were calculated with mothur software and plotted on a NMDS graph. Samples from different individuals of the same species are represented by the same color.

Seawater samples showed greater microbiota diversity (Fig 7, Fig S3) than sponges, and the sample from Fields Bay, Antarctica, was the least diverse among them. Among the sponge species, *P. magna* presented the lowest Shannon diversity index, while all other samples presented higher and similar values (Fig 7).

**Figure 7.**
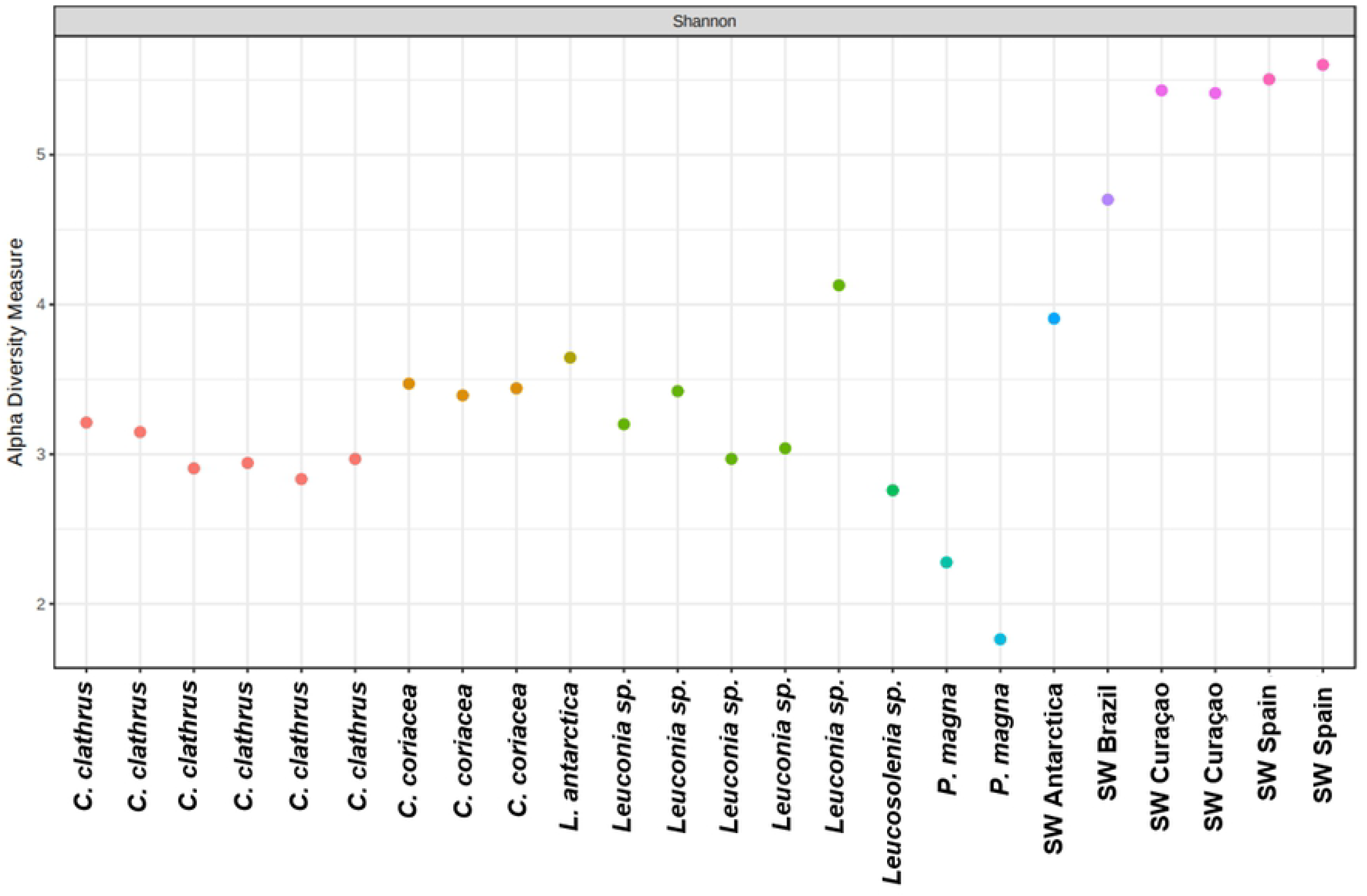
Alpha diversity analysis of microbiota from different calcareous sponge species and seawater samples. The dispersion plot was built based on Shannon index of diversity.

A functional pathway comparison of the microbiomes predicted with PICRUSt software shows that, although the taxonomic composition of the microbiota is different, there is a level of conservation in which metabolic features are present in sponge and seawater samples, preserving the specificity of each symbiotic relationship, as the different species and seawater samples cluster together on a hierarchical clustering cladogram based on Bray-Curtis dissimilarity (Fig S4). Heatmap analysis of the 15 most abundant pathways indicates prevalence, in all samples, of heterotrophic metabolism and nutrient transport with higher incidence of carbohydrate and amino acids metabolism (Fig 8). Cellular housekeeping functions, nucleotide and lipid metabolism were the least abundant functions. Statistical analysis shows that pathways related to cell motility, membrane transport, genetic information processing, xenobiotics metabolism and signal transduction are higher in sponges (Fig 9). Meanwhile, amino acid and nucleotide metabolism, translation, replication and repair, folding, sorting and degradation and glycan biosynthesis and metabolism were more abundant in seawater symbionts.

**Figure 8.**
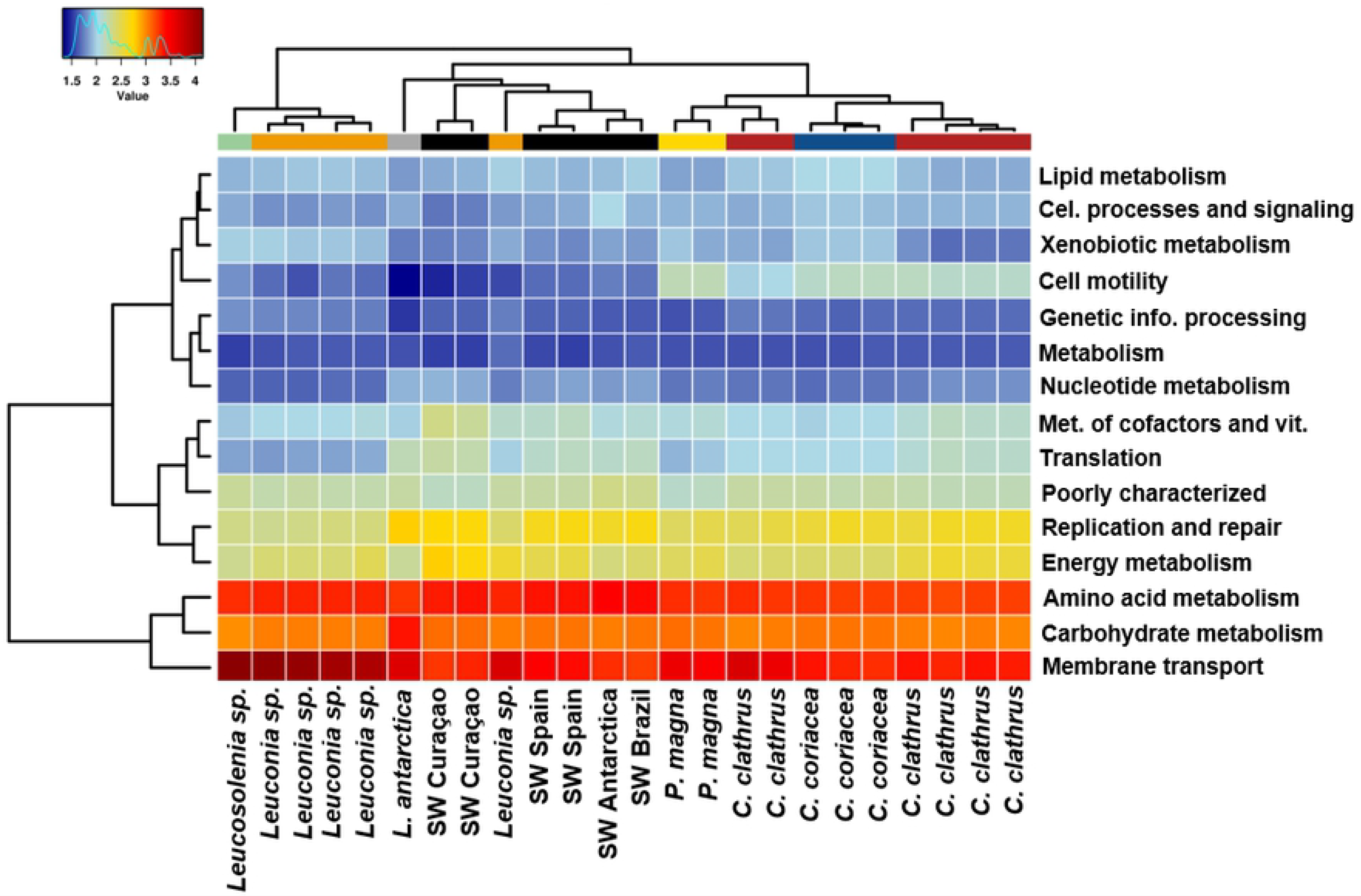
Abundance heatmap of predicted metagenomic functional profiles from different calcareous sponge species and seawater samples. Heatmap shows the 15 most abundant metabolic pathways predicted with PICRUSt. Colors shift from blue (lower) to red (higher) according to pathway abundance in each sample.

**Figure 9.**
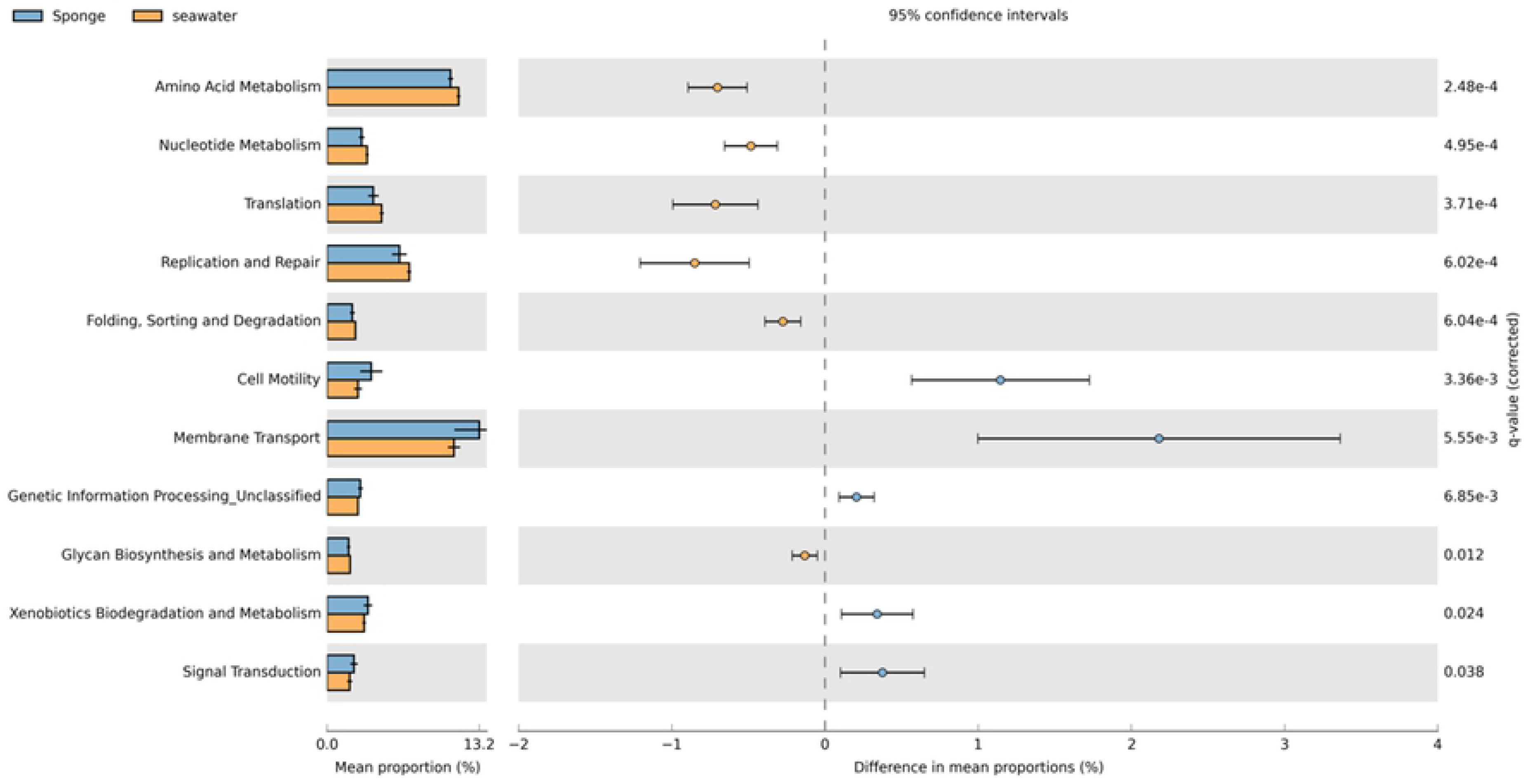
Statistical analysis of the differential abundance of predicted metagenomic functional profiles from sponges and seawater samples. Statistical analysis of the predicted metagenomes was performed with STAMP software.

## Discussion

In 2017, Moitinho-Silva *et al.* [7] described a worldwide initiative to study sponge microbiomes. It was reported that on the Earth Microbiome Project (EMP) [35] database there is sequencing data for over three thousand samples, representing 269 sponge species from all over the world. However, no data was present for sponges of the Brazilian coast. In addition, the microbiota of calcareous sponges has been understudied despite the fact that initial investigations using cultivation procedures indicate that these animals are a potential source of microbial biodiversity and of bioactive molecules [36–38].

The study reported here presents the first description of *P. magna* microbiota by next generation sequencing and increases our knowledge of the microbial symbionts of calcareous sponges that inhabit the Brazilian coast. We observed that *P. magna* microbiota has low diversity, as indicated by the inverse Simpson index, and by the observation that only two OTUs corresponded to 80% (*P. magna* 6) and 90% (*P. magna* 8) of all sequences. Meanwhile, all other OTUs were not represented above 0.05%. These rare OTUs might even be residue from the feeding habits of these animals. Giles *et al*. [39] previously described this microbial profile, of few OTUs with high relative abundance, on LMA sponges.

*Paraleucilla magna*’s microbiota is composed predominantly of microorganisms from the Proteobacteria phylum, following the pattern observed for LMA sponges (Fig 2). The two most abundant OTUs could not be assigned to any known genus. OTU001 could only be classified in the class Alphaproteobacteria and OTU002 was allocated in the family Rhodospirillaceae. The Alphaproteobacteria are a quite diverse group of microorganisms with important biological roles [40]. In the phylogenetic tree, OTU001 sequence did not cluster closely with any related sequence. The closest similarity was low and with a quite diverse group of microorganisms detected in various locations, being two from deep-sea regions, one from *Paeonia ostii*’s rhizosphere and one from a hyper saline mat (Fig 3A).

The family Rhodospirillaceae, to which OTU002 was assigned, has 34 known genera of Gram-negative microorganisms that present various nutritional strategies [41]. In the phylogenetic tree constructed with related sequences retrieved from GenBank, OTU002 showed similarity with marine bacteria from different habitats, grouping closely with a sequence from a bacterium isolated from a marine biofilm.

Yarza *et al.* [42] reported a phenomenon in which, as sequencing and data processing technologies develop, the molecular description of new taxa extrapolate the classical taxonomy system, leaving a great number of microorganisms with no classification. This seems to be the case for OTU002. Differently, OTU001 appears to have not been sequenced yet, as the highest similarity value with other reported sequences in Genbank was 91%.

The seawater sample showed the greatest microbial diversity, and 35% of the OTUs did not reach a relative abundance of at least 5%. The most abundant OTU belongs to the SAR92 clade, a member of the Gammaproteobacteria (Fig 2). This group comprehends oligotrophic marine organisms, capable of producing proteorhodopsin, an enzyme that enables the generation of energy from sunlight. This allows a nutritional optimization in environments where carbon concentration is low. This clade is often found in coastal phototrophic zones, such as where our sample was collected [43].

The predicted bacterial metagenomes for *P. magna* individuals (Fig 5A) revealed that the most abundant functional pathways were related to carbohydrate and amino acid metabolism and membrane transport, indicating the participation of the symbionts in the obtention and transport of nutrients. The seawater predicted metagenome was distinguished mainly for having a lower abundance in pathways related to nutrient metabolism, which corroborates the fact that the most abundant genera in this sample are oligotrophic.

Karimi *et al.* [44] published a study reporting the main genomic features of alphaproteobacterial sponge symbionts. Corroborating our data, their study showed a prevalence of ABC transporters, fundamental for nutrient obtention, and versatile nutrients metabolism, pointing to a great importance in nutrient cycling. Moreover, they also described a reduction in motility observed in symbionts, contrary to that observed in free-living bacteria. In our heatmap (Fig 5), we also observed a low abundance in motility pathways.

In the domain Archaea, the phylum Thaumarcheota was predominant in *P. magna* samples, as already described by studies with other sponges (Fig 4) [45]. On a lower taxonomic rank, the genus candidatus *Nitrosopumillus* was the most abundant, followed by *Nitrosopellagicus.* In the seawater sample an almost equal proportion of Thaumarcheota and Euryarcheota were found. Accordingly, a previous study performed in two sites at Guanabara bay in Rio de Janeiro, using clone libraries of the archaeal ammonia oxidase gene, *amoA*, reported the absence of Euryarcheota and the presence of a distinctive community of Thaumarcheota in *P. magna* samples [46].

Few studies have been performed using archaeal specific primers, however, a consistency was found between the findings using this methodology and 16S rRNA sequencing. Our approach used the same primer for archaeal and bacterial kingdoms, leading to a possible loss of resolution in the identification of the former. In spite of this possibility, no studies have reported limitation due to this choice of primers in the literature [47,48].

The phylum Thaumarcheota and the two genera found in our samples show metabolic activity related to the nitrogen biogeochemical cycle, standing out as an essential group for nitrification activities, using the generated energy for CO_2_ fixation [45]. Also, this group has an important role as producer of cobalamine, an essential co-factor for animal life. As such, the predicted archaeal metagenomes of *P. magna* samples show a high abundance of pathways related to the metabolism of nitrogenated compounds and energy metabolism. Other functional categories with high abundance were those related to cellular maintenance.

Euryarcheota genomic data indicate participation in the degradation of proteins, lipids and vitamins [49,50]. The predicted archaeal metagenome for seawater, presents itself as a combination of the two observed phyla, with high abundance of nucleotide and carbohydrate metabolisms, showing the importance of these two phyla for nutrient cycling in the ocean.

Data shown here indicate that *P. magna* is most likely a LMA sponge, as it presents a microbiota dominated by one phylum, represented by few and abundant OTUs. When a large number of sponges from the class Demospongiae were analyzed, researchers were not able to find a prokaryotic signature of the class, as the reported microbiota compositions were usually species-specific [51]. In order to investigate this pattern in the class Calcarea, we analyzed the available microbiota found for other species available in the SRA database. We were able to obtain bacterial V4 16S rRNA sequence data from five species: *Clathrina clathrus*, *C. coriacea* and *Leucetta antarctica*, from the subclass Calcinea and *Leucosolenia* sp. and *Leuconia* sp. from the subclass Calcaronea. We also analyzed the microbiota from four different seawater samples to compare the microbiota of the sponges and the environment they inhabit, which is fundamental for LMA sponge studies.

The microbiota of all sponge species was clearly separated from those of seawater samples both on the NMDS plot (Fig 6) and in the dendogram (Fig S2). Seawater samples clustered together, preserving certain dissimilarities. Among the sponges, it has been shown that short phylogenetic distances correspond to more similar microbiota [52,53]. In the present study, we could observe this phenomenon in two distinct species of the same genus, *Clathrina clathrus* and *C. coriacea*, clustered together in the NMDS plot (Fig 6) and on the cladogram (Fig S2). These two sponges species are of the same subclass as *L. antarctica*, however, no similarity in microbiota composition was found, with the latter forming a separate branch on the cladogram. The species *Leucosolenia* sp. and *P. magna* are of the same order, Leucosolenida, but they did not cluster together. In fact, *Leucosolenia* sp., a species of the subclass Calcaronea, clustered with species of *Clathrina*, i.e. species of the subclass Calcinea (Fig S2). Although they are phylogenetically distant, they have the same type of aquiferous system, asconoid. *Paraleucilla* and *Leuconia* samples also formed a cluster (Fig S2) and are in proximity on the NMDS graph (Fig 6). Both species have a leuconoid aquiferous system.

Although the microbiota of only a few calcareous sponges has been studied, the clustering of asconoid and leuconoid species is very interesting. In the asconoid aquiferous system, the body wall of the sponge is very thin, perhaps allowing higher concentrations of O_2_ in the mesohyl. Moreover, this thin body wall allows more light penetration. On the other hand, leuconoid sponges have thicker body walls, what may result in less O_2_ in the mesohyl, albeit the presence of many canals and choanocytary chambers. Moreover, there is lesser light penetration in the mesohyl. Both O_2_ concentrations and light penetration could be related to microorganism selection, explaining the proximity of sponges with the same aquiferous system.

Microbiota composition of the seawater samples was more diverse than those of the sponge species. The seawater sample from Bay Fields, Antarctica, was the least diverse and those from Montoya, Spain, showed the highest diversity. *P. magna* is the sponge species with the lowest microbial diversity, while *C. coriacea* and *L. antarctica* are the most diverse, based on the Shannon index (Fig 7). This index calculates diversity based on richness and abundance of microorganisms combined and although no index has been deemed as ideal, a study reported the Shannon index as being capable of describing the largest number of relationships/traits [54].

Hierarchical clustering based on the predicted bacterial metabolism showed that sponges from the same genus share similar metabolic pathways and these are clearly separate from seawater samples, indicating that there is a fundamental difference between the microorganisms inhabiting these two habitats and the functions they perform (Fig S4). This can also be seen in the heatmap of the 15 most abundant predicted pathways and in the statistical analysis with STAMP (Fig 8 and Fig 9, respectively), in which functions related to nutrient acquisition (membrane transport, xenobiotics biodegradation and metabolism) and symbiont/host interactions (cell motility, genetic information processing and signal transduction) were prevalent in sponges and functions related to cellular metabolism (amino acid metabolism) and proliferation (nucleotide metabolism, replication and repair) had higher abundance in seawater (Fig 9). Conversely, Karimi *et al.* [55], showed that ABC transporters were more prevalent in sponges.

This study shows the first description of *P. magna*’s microbiota by next-generation sequencing. This microbiota is characteristic of an LMA sponge, is dominated by few Alphaproteobacteria OTUs and has a predicted metabolism directed to nutrient uptake and degradation and housekeeping functions. Also, when compared to other species of calcareous sponges, *P. magna* symbionts differed at both OTU and metabolic levels. Other studies need to be performed in order to determine if *P. magna* presents a stable microbiota across seasonal and geographical distances.

## Acknowledgments

This work was partially supported by grants from Fundação Carlos Chagas Filho de Amparo à Pesquisa do Estado do Rio de Janeiro, Conselho Nacional de Desenvolvimento Científico e Tecnológico and Coordenação de Aperfeiçoamento de Pessoal de Nível Superior – Brasil (CAPES) – Finance Code 001.

## Supporting information captions

**Figure S1. Rarefaction analysis of microbiota from *P. magna* and seawater samples**. (A) Bacteria domain and (B) Archaea domain.

**Figure S2. Hierarchical clustering of microbiota composition from different calcareous sponge species and seawater samples.** Distance among groups was defined by the Yue & Clayton theta calculator with mothur software.

**Figure S3. Rarefaction analysis of microbial community richness of different calcareous sponge species and seawater samples.**

**Figure S4. Hierarchical clustering according to Bray-Curtis distances of predicted functional profiles from different calcareous sponge species and seawater samples.** Metabolic pathways were predicted with PICRUSt and Bray-Curtis distances were calculated with Calypso software.

## References

1. van Soest RWM, Boury-Esnault N, Vacelet J, Dohrmann M, Erpenbeck D, de Voogd NJ, et al. Global diversity of sponges (Porifera). PLoS One. 2012;7. doi:10.1371/journal.pone.0035105

2. Hentschel U, Piel J, Degnan SM, Taylor MW. Genomic insights into the marine sponge microbiome. Nat Rev Microbiol. Nature Publishing Group; 2012;10: 641–654. doi:10.1038/nrmicro2839

3. Webster NS, Thomas T. The sponge hologenome. MBio. 2016;7: 1–14. doi:10.1128/mBio.00135-16

4. Gloeckner V, Wehrl M, Moitinho-Silva L, Gernert C, Hentschel U, Schupp P, et al. The HMA-LMA dichotomy revisited: An electron microscopical survey of 56 sponge species. Biol Bull. 2014;227: 78–88. doi:10.1086/BBLv227n1p78

5. Abdelmohsen UR, Balasubramanian S, Oelschlaeger TA, Grkovic T, Pham NB, Quinn RJ, et al. Potential of marine natural products against drug-resistant fungal, viral, and parasitic infections. Lancet Infect Dis. Elsevier Ltd; 2017;17: e30–e41. doi:10.1016/S1473-3099(16)30323-1

6. Thomas T, Moitinho-Silva L, Lurgi M, Björk JR, Easson C, Astudillo-García C, et al. Diversity, structure and convergent evolution of the global sponge microbiome. Nat Commun. 2016;7. doi:10.1038/ncomms11870

7. Moitinho-Silva L, Nielsen S, Amir A, Gonzalez A, Ackermann GL, Cerrano C, et al. The sponge microbiome project. Gigascience. 2017;6: 1–7. doi:10.1093/gigascience/gix077

8. Rodríguez-Marconi S, De La Iglesia R, Díez B, Fonseca CA, Hajdu E, Trefault N, et al. Characterization of bacterial, archaeal and eukaryote symbionts from antarctic sponges reveals a high diversity at a three-domain level and a particular signature for this ecosystem. PLoS One. 2015;10: 1–19. doi:10.1371/journal.pone.0138837

9. Klautau M, Monteiro L, Borojevic R. First occurrence of the genus Paraleucilla (Calcarea, Porifera) in the Atlantic Ocean: P. magna sp nov. Zootaxa. 2004;8: 1–8. doi:10.5281/zenodo.158320

10. Longo C, Mastrototaro F, Corriero G. Occurrence of Paraleucilla magna (Porifera: Calcarea) in the Mediterranean Sea. J Mar Biol Assoc UK. Cambridge University Press; 2007;87: 1749–1755. doi:10.1017/S0025315407057748

11. Guardiola M, Frotscher J, María •, Uriz J. Genetic structure and differentiation at a short-time scale of the introduced calcarean sponge Paraleucilla magna to the western Mediterranean. doi:10.1007/s10750-011-0948-1

12. Biology M, Biology M. New data on the distribution of the alien sponge Paraleucilla magna Klautau, Monteiro & Borojevic, 2004 in the … sponge Paraleucilla magna Klautau,. 2016;

13. Agell G, Frotscher J, Guardiola M, Pascual M, Uriz MJ. Characterization of nine polymorphic microsatellite loci for the calcareous sponge Paraleucilla magna Klautau et al. 2004 introduced to the Mediterranean Sea. Conserv Genet Resour. 2012;4: 403–405. doi:10.1007/s12686-011-9560-y

14. Padua A, Lanna E, Klautau M. Macrofauna inhabiting the sponge Paraleucilla magna (Porifera: Calcarea) in Rio de Janeiro, Brazil. doi:10.1017/S0025315412001804

15. Santos OCS, Pontes PVML, Santos JFM, Muricy G, Giambiagi-deMarval M, Laport MS. Isolation, characterization and phylogeny of sponge-associated bacteria with antimicrobial activities from Brazil. Res Microbiol. Elsevier Masson SAS; 2010;161: 604–612. doi:10.1016/j.resmic.2010.05.013

16. Lanna E, Klautau M. Embryogenesis and larval ultrastructure in Paraleucilla magna (Calcarea, Calcaronea), with remarks on the epilarval trophocyte epithelium (“placental membrane”). Zoomorphology. Springer-Verlag; 2012;131: 277–292. doi:10.1007/s00435-012-0160-5

17. Lanna E, Paranhos R, Paiva PC, Klautau M. Environmental effects on the reproduction and fecundity of the introduced calcareous sponge Paraleucilla magna in Rio de Janeiro, Brazil. Mar Ecol. Wiley/Blackwell (10.1111); 2015;36: 1075–1087. doi:10.1111/maec.12202

18. Redmond NE, van Soest RWM, Kelly M, Raleigh J, Travers SAA, McCormack GP. Reassessment of the classification of the Order Haplosclerida (Class Demospongiae, Phylum Porifera) using 18S rRNA gene sequence data. Mol Phylogenet Evol. 2007;43: 344–352. doi:10.1016/j.ympev.2006.10.021

19. Altschul SF, Gish W, Miller W, Myers EW, Lipman DJ. Basic local alignment search tool. J Mol Biol. Academic Press; 1990;215: 403–410. doi:10.1016/S0022-2836(05)80360-2

20. Benson, Dennis; Karsch-Mizrachi, Ilene; Ostel, James; Wheeler D. Genbank. Nucleic Acids Res. 2011;39.

21. Katoh K, Standley DM. Article Fast Track MAFFT Multiple Sequence Alignment Software Version 7: Improvements in Performance and Usability. doi:10.1093/molbev/mst010

22. Kumar S, Stecher G, Tamura K, Dudley J. MEGA7: Molecular Evolutionary Genetics Analysis Version 7.0 for Bigger Datasets. Mol Biol Evol. 2016;33: 1870–1874. doi:10.1093/molbev/msw054

23. Caporaso JG, Lauber CL, Walters WA, Berg-Lyons D, Lozupone CA, Turnbaugh PJ, et al. Global patterns of 16S rRNA diversity at a depth of millions of sequences per sample. Proc Natl Acad Sci. 2011;108: 4516–4522. doi:10.1073/pnas.1000080107

24. Schloss PD, Westcott SL, Ryabin T, Hall JR, Hartmann M, Hollister EB, et al. Introducing mothur: Open-Source, Platform-Independent, Community-Supported Software for Describing and Comparing Microbial Communities. Appl Environ Microbiol. 2009;75: 7537–7541. doi:10.1128/AEM.01541-09

25. Kozich JJ, Westcott SL, Baxter NT, Highlander SK, Schloss PD. Development of a dual-index sequencing strategy and curation pipeline for analyzing amplicon sequence data on the miseq illumina sequencing platform. Appl Environ Microbiol. 2013;79: 5112–5120. doi:10.1128/AEM.01043-13

26. Quast C, Pruesse E, Yilmaz P, Gerken J, Schweer T, Yarza P, et al. The SILVA ribosomal RNA gene database project: improved data processing and web-based tools. Nucleic Acids Res. Oxford University Press; 2012;41: D590–D596. doi:10.1093/nar/gks1219

27. Rognes T, Flouri T, Nichols B, Quince C, Mahé F. VSEARCH: a versatile open source tool for metagenomics. PeerJ. PeerJ Inc.; 2016;4: e2584. doi:10.7717/peerj.2584

28. Dhariwal A, Chong J, Habib S, King IL, Agellon LB, Xia J. MicrobiomeAnalyst: a web-based tool for comprehensive statistical, visual and meta-analysis of microbiome data. Nucleic Acids Res. Oxford University Press; 2017;45: W180–W188. doi:10.1093/nar/gkx295

29. DeSantis TZ, Hugenholtz P, Larsen N, Rojas M, Brodie EL, Keller K, et al. Greengenes, a chimera-checked 16S rRNA gene database and workbench compatible with ARB. Appl Environ Microbiol. American Society for Microbiology; 2006;72: 5069–72. doi:10.1128/AEM.03006-05

30. Langille MGI, Zaneveld J, Caporaso JG, McDonald D, Knights D, Reyes JA, et al. Predictive functional profiling of microbial communities using 16S rRNA marker gene sequences. Nat Biotechnol. 2013;31: 814–821. doi:10.1038/nbt.2676

31. Kanehisa M, Sato Y, Kawashima M, Furumichi M, Tanabe M. KEGG as a reference resource for gene and protein annotation. Nucleic Acids Res. 2015;44: 457–462. doi:10.1093/nar/gkv1070

32. Zakrzewski M, Proietti C, Ellis JJ, Hasan S, Brion M-J, Berger B, et al. Calypso: a user-friendly web-server for mining and visualizing microbiome–environment interactions. Bioinformatics. Oxford University Press; 2016;33: btw725. doi:10.1093/bioinformatics/btw725

33. Parks D, Beiko R. STAMP User’s Guide v2. 2011; Available: http://kiwi.cs.dal.ca/Software/images/0/02/STAMP_Users_Guide_v2.0.0.pdf

34. Hirayama H, Sunamura M, Takai K, Nunoura T, Noguchi T, Oida H, et al. Culture-Dependent and-Independent Characterization of Microbial Communities Associated with a Shallow Submarine Hydrothermal System Occurring within a Coral Reef off Taketomi Island, Japan. Appl Environ Microbiol. 2007;73: 7642–7656. doi:10.1128/AEM.01258-07

35. Thompson LR, Sanders JG, McDonald D, Amir A, Ladau J, Locey KJ, et al. A communal catalogue reveals Earth’s multiscale microbial diversity. Nature. 2017;551: 457–463. doi:10.1038/nature24621

36. Fromont, Jane; Huggett, Megan; Lengger, Sabine; Grice, Kitti; Schonberg C. Characterization of Leucetta prolifera, a calcarean cyanosponge from south-western Australia, and its symbionts. J Mar Biol Assoc United Kingdom. 2014; doi:10.1111/j.1462-2920.2008.01815.x

37. Roué M, Quévrain E, Domart-Coulon I, Bourguet-Kondracki ML. Assessing calcareous sponges and their associated bacteria for the discovery of new bioactive natural products. Nat Prod Rep. 2012;29: 739–751. doi:10.1039/c2np20040f

38. Flemer B, Kennedy J, Margassery LM, Morrissey JP, O’Gara F, Dobson ADW. Diversity and antimicrobial activities of microbes from two Irish marine sponges, Suberites carnosus and Leucosolenia sp. J Appl Microbiol. 2012;112: 289–301. doi:10.1111/j.1365-2672.2011.05211.x

39. Giles EC, Kamke J, Moitinho-Silva L, Taylor MW, Hentschel U, Ravasi T, et al. Bacterial community profiles in low microbial abundance sponges. FEMS Microbiol Ecol. 2013;83: 232–241. doi:10.1111/j.1574-6941.2012.01467.x

40. Williams KP, Sobral BW, Dickerman AW. A Robust Species Tree for the Alphaproteobacteria † Downloaded from. J Bacteriol. 2007;189: 4578–4586. doi:10.1128/JB.00269-07

41. Baldani JI, Videira SS, dos Santos Teixeira KR, Reis VM, de Oliveira ALM, Schwab S, et al. The Family Rhodospirillaceae. The Prokaryotes. Berlin, Heidelberg: Springer Berlin Heidelberg; 2014. pp. 533–618. doi:10.1007/978-3-642-30197-1_300

42. Yarza P, Yilmaz P, Pruesse E, Glöckner FO, Ludwig W, Schleifer K-H, et al. Uniting the classification of cultured and uncultured bacteria and archaea using 16S rRNA gene sequences. Nat Rev Microbiol. Nature Publishing Group; 2014;12: 635–645. doi:10.1038/nrmicro3330

43. Stingl U, Desiderio RA, Cho J-C, Vergin KL, Giovannoni SJ. The SAR92 clade: An abundant coastal clade of culturable marine bacteria possessing proteorhodopsin A C C E P T E D. Appl Environ Microbiol. 2007; doi:10.1128/AEM.02559-06

44. Karimi E, Slaby BM, Soares AR, Blom J, Hentschel U, Costa R. Metagenomic binning reveals versatile nutrient cycling and distinct adaptive features in alphaproteobacterial symbionts of marine sponges. FEMS Microbiol Ecol. Oxford University Press; 2018;94. doi:10.1093/femsec/fiy074

45. Zhang F, Pita L, Erwin PM, Abaid S, López-Legentil S, Hill RT. Symbiotic archaea in marine sponges show stability and host specificity in community structure and ammonia oxidation functionality. FEMS Microbiol Ecol. Oxford University Press; 2014;90: 699–707. doi:10.1111/1574-6941.12427

46. Turque AS, Batista D, Silveira CB, Cardoso AM, Vieira RP, Moraes FC, et al. Environmental shaping of sponge associated archaeal communities. PLoS One. 2010;5. doi:10.1371/journal.pone.0015774

47. Brochier-Armanet C, Boussau B, Gribaldo S, Forterre P. Mesophilic crenarchaeota: proposal for a third archaeal phylum, the Thaumarchaeota. Nat Rev Microbiol. Nature Publishing Group; 2008;6: 245–252. doi:10.1038/nrmicro1852

48. Pester M, Wagner M. The Thaumarchaeota: an emerging view of their phylogeny and ecophysiology. Curr Opin Microbiol. Elsevier Current Trends; 2011;14: 300–306. doi:10.1016/J.MIB.2011.04.007

49. Iverson V, Morris RM, Frazar CD, Berthiaume CT, Morales RL, Armbrust EV. Untangling Genomes from Metagenomes: Revealing an Uncultured Class of Marine Euryarchaeota. doi:10.1126/science.1093857

50. Zhang CL, Xie W, Martin-Cuadrado A-B, Rodriguez-Valera F. Marine Group II Archaea, potentially important players in the global ocean carbon cycle. Front Microbiol. Frontiers; 2015;6: 1108. doi:10.3389/fmicb.2015.01108

51. Steinert G, Rohde S, Janussen D, Blaurock C, Schupp PJ. Host-specific assembly of sponge-associated prokaryotes at high taxonomic ranks OPEN. doi:10.1038/s41598-017-02656-6

52. Easson CG, Thacker RW. Phylogenetic signal in the community structure of host-specific microbiomes of tropical marine sponges. Front Microbiol. 2014;5: 1–11. doi:10.3389/fmicb.2014.00532

53. Souza DT, Genuário DB, Silva FSP, Pansa CC, Kavamura VN, Moraes FC, et al. Analysis of bacterial composition in marine sponges reveals the influence of host phylogeny and environment. Olson J, editor. FEMS Microbiol Ecol. Oxford University Press; 2017;93: fiw204. doi:10.1093/femsec/fiw204

54. Morris EK, Caruso T, Buscot F, Fischer M, Hancock C, Maier TS, et al. Choosing and using diversity indices: insights for ecological applications from the German Biodiversity Exploratories. Ecol Evol. Wiley-Blackwell; 2014;4: 3514–3524. doi:10.1002/ece3.1155

55. R. EKMRJGJXMR. Comparative Metagenomics Reveals the Distinctive Adaptive Features of the Spongia officinalis Endosymbiotic Consortium. 2017; doi:10.3389/fmicb.2017.02499

